# Functional Profiling of Thousands of Sequence-Diverse Protease Homologs with GROQ-seq

**DOI:** 10.64898/2026.04.30.721934

**Authors:** James R. McLellan, Svetlana Ikonomova, Shwetha Sreenivasan, Alan N. Amin, Catherine Baranowski, Amanda Reider Apel, Peter Kelly, David Ross, Aviv Spinner

## Abstract

High-quality datasets that span broad sequence diversity are essential for understanding protein sequence–function relationships beyond local mutational landscapes. Here, we applied Growth-based Quantitative Sequencing (GROQ-seq) to measure function across an 11,722 member protease library, comprised of natural homologs and AI-shrunken variants. This library spans vast sequence diversity, with Levenshtein distances of up to 245 and a mean pairwise sequence identity of 41% to TEV protease S219V. We identified sequence-divergent TEV protease homologs that preserve function against the native TEV protease substrate. These findings reveal the robustness of protease activity across highly diverse sequences. Here, we demonstrate the aptitude of the GROQ-seq assay for screening large, diverse protein libraries for function, enabling efficient data generation at scale for training machine learning models across broad sequence landscapes.

## Introduction

Protein language models rely on large, sequence-diverse training datasets that map sequence to function.^1^ Therefore, to enable machine learning models that accurately predict protein function from sequence, AI-ready biological datasets must deliver functional measurements of large and diverse protein libraries. Commonly used approaches build comprehensive mutagenesis libraries from a single reference sequence.^2^ However, homologous proteins can often perform the same function through similar structural motifs despite highly dissimilar sequences.^3^ Prior studies have generated mutagenesis datasets across phylogenetically-diverse libraries, including dihydrofolate reductase (DHFR), nuclease enzymes, and DNA binding proteins, with approaches including homolog library construction, machine learning-guided design, and ancestral inference, to map the sequence-function landscape across tens of thousands of variants.^4–7^ To efficiently produce datasets for sequence-diverse proteins, there is a need to explore more broad mutational scanning (BMS) approaches, in which libraries and assays connect sequence, structure, and function across a wide breadth of sequences.

Previously, we employed the GROQ-seq assay (**Supplementary Fig. 1**) to measure the function of 3 different protein classes: transcriptional repressors, an RNA polymerase, and the Tobacco Etch Virus (TEV) protease. Using site-saturation variant libraries (SSVL) and error-prone PCR (epPCR), we quantified mutational effects across tens of thousands of variants and mapped their sequence-function landscapes.^8^

Here, we applied GROQ-seq to characterize libraries spanning a vastly divergent evolutionary sequence space. In this report, we present a dataset that captures the sequence-function relationships across a sequence-diverse set of TEV protease natural homologs and AI-generated variants. We quantified their function against the canonical TEV protease cleavage site using GROQ-seq (see reproducibility report for assay details).^9^

This dataset demonstrates that protease function can persist despite high sequence diversity, highlighting the flexibility of the sequence-function relationship. We identified active homologs with as little as 19% sequence identity to TEV protease, as well as shrunken variants, with up to 70 residues removed, which retain measurable activity. This work underscores the breadth of sequence variation that can preserve protease function and demonstrates the utility of the GROQ-seq assay for analyzing sequence-diverse protein libraries at scale.

## Methods

### Protein variant library design

Using TEV protease S219V (a commonly used mutant of TEV wild-type in biotechnology research to avoid autoproteolysis^10^) as our starting reference, we performed five iterations of JackHMMER^11^ within the EVCouplings^12^ pipeline to retrieve TEV protease natural homolog sequences on UniProt100^13^, BFD^14^, and MGnify^15^. Because the initial alignments frequently lacked complete N- and C-terminal coverage, we retrieved full-length sequences from the source databases using the corresponding sequence identifiers. As these records often represented full viral genomes, we individually aligned each sequence to the TEV reference to extract the full-length protease region, which we used as the homologous sequences in this study. Using this approach, we retrieved a total of 4,022 TEV protease homologs. Additionally, we used SCISOR, a discrete diffusion model trained with a denoising task that learns to remove randomly inserted residues, to generate shortened, “shrunken,” variants of the TEV protease.^16^ In this context, “shrunken” denotes the deletion of residues throughout the sequence (not restricted to contiguous regions).

We included 422 shrunken variants with up to 150 amino acid deletions in the library design for this dataset. We then codon optimized each sequence for expression in *Escherichia coli* using DNA Chisel and avoiding restriction sites for BsaI, BsmBI, SapI, BspQI, BtsI, and PaqCI.^17^

### Variant library generation

The 4,444 sequences in the designed library were synthesized by SynPlexity (https://synplexity.com). Synthesis of full-length genes introduced a degree of additional random variation into the library, which expanded the number of variants to 11,722 unique amino acid sequences. The variants and random barcodes were cloned into the plasmid backbone via Golden Gate Assembly by Clutch Biotechnologies (Burlingame, California). Oxford Nanopore Technology performed by Plasmidsaurus (Plasmidsaurus Inc., USA) was used to map barcodes to variants. Libraries were transformed into the *E. coli* Marionette-Clo strain.^18^ Summary statistics of our final library build can be seen in **Table 1**.

**Table 1.** Summary statistics of TEV protease homolog library. Pairwise identity for the set of sequences in the final library was calculated from the multiple sequence alignment made using MAFFT’s progressive algorithm (FFT-NS-2).^19^.

| No. Natural Homologs | No. Shrunken TEV Proteases | Final No. Unique Sequences | Minimum Sequence Pairwise Identity | Median Sequence Pairwise Identity | Mean Sequence Pairwise Identity |
| --- | --- | --- | --- | --- | --- |
| 4,022 | 422 | 11,722 | 6.52% | 50.23% | 41.79% |

### Circuit design and GROQ-seq assay

The plasmid backbone for this circuit was previously described.^8^ Briefly, we cloned each variant into a plasmid backbone containing expression of a split-DHFR protein, where two DHFR fragments are linked by the TEV protease substrate (ENLYFQS) (**Supplementary Fig. 2**). This links protease function to cellular fitness under trimethoprim (TMP) stress.^8^ Proteases with catalytic activity for that substrate cleave the split-DHFR, thereby reducing cellular fitness in the presence of TMP. Cells which contain low-functioning variants will not cleave DHFR and maintain resistance to TMP, resulting in higher cellular fitness. We included a calibration ladder in each sample to convert from cellular fitness (measured via DNA barcode sequencing) to the level of intact DHFR substrate in order to give a more biophysically meaningful result.^8,20^ We also implemented the GROQ-seq protease assay in a multi-parametric format, with measurements of intact DHFR at 10 different expression levels of the protease and 2 different expression levels of the DHFR substrate. For simplicity, in this report, we assess the function of TEV protease homologs using the assay condition with the highest level of protease expression and a low DHFR expression, which was previously demonstrated to yield the highest dynamic range of protease function in the assay.^8^ Function measurements are reported as the base-10 logarithm of the level of intact DHFR, with lower levels of intact DHFR corresponding to higher protease activity.

### Data Availability

The GROQ-seq Fitness and Function Measurements for TEV Protease Homolog Library dataset used in this report can be accessed from the DOI: 10.5281/zenodo.19926758.^21^

## Results

### TEV Homolog and Shrunk TEV Library Samples a Highly Diverse Sequence Space

We compared the sequence diversity landscape of our natural homolog library with respect to TEV S219V. To this end, we made a multiple sequence alignment (MSA) of the 11,722 sequences using MAFFT’s progressive algorithm (FFT-NS-2). This alignment (without any redundancy filtering) was used to identify 233 match columns with <50% gaps across all sequences. This culled version of the MSA was used to construct a maximum-likelihood phylogenetic tree for our library using FastTree 2.1.11 (LG substitution model, Gamma20 rate variation, four SPR refinement rounds) (**Fig. 1A**).

**Figure 1.**
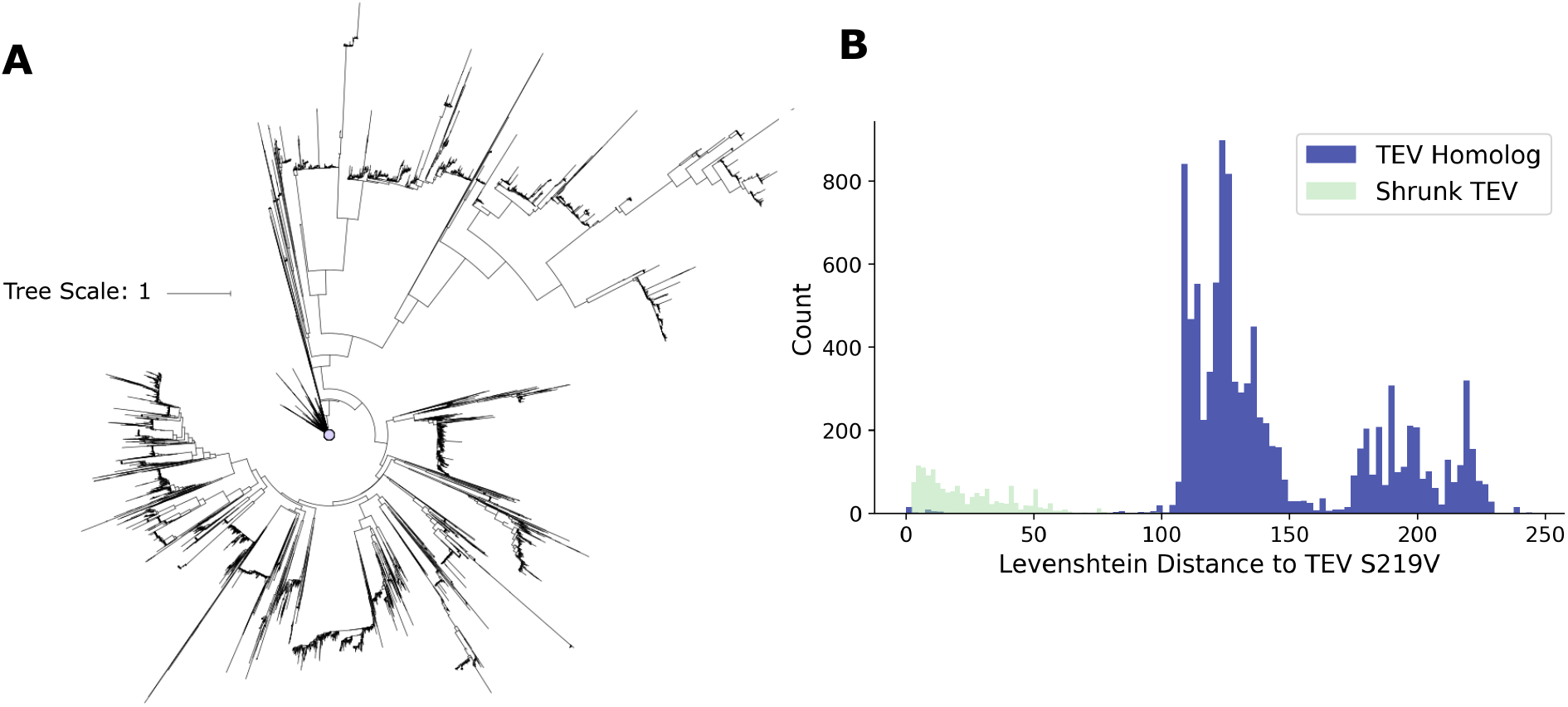
TEV Homolog Sequence Diversity **A.** Maximum-likelihood tree of 11,722 protease homologs and shrunken variants in this report inferred using FastTree 2.1.11 (LG substitution model) from MAFFT multiple sequence alignment restricted to 233 match columns with <50% gaps across all sequences). The reference TEV S219V sequence used in this report is marked with the lavender circle at the top of the tree. **B**. Levenshtein distance distribution for homolog and shrunken variants relative to TEV S219V.

Notably, TEV S219V is positioned at the top of the tree, within the densely populated center (**Fig. 1A**). Near TEV S219V, densely clustered clades of closely related homologs extend from short branches. Distal clades arising from elongated branches represent greater leaps in sequence divergence, yet distinct clades are still observed at the terminal ends.

To further quantify the phylogenetic diversity of the library, we computed the pairwise identity matrix and examined the distribution of all sequences across different identity thresholds with respect to TEV S219V (**Supplementary Table 1**). Only 13% of the library shares ≥ 70% identity with TEV S219V, and only 56% of the library shares ≥ 50% identity. This indicates that nearly half the library sits below the conventional threshold for functional conservation.^22^ Next, we computed the effective number of sequences (N_eff) at different identity thresholds using the standard sequence-reweighting approach (**Supplementary Table 1**).^23^ At 80% identity threshold, the library of 11,722 sequences collapses to 301 effective clusters. This reiterates the observation that a small, dense group of homologs appear near TEV S219V and a broad tail extending beyond. Furthermore, the pairwise identity to TEV S219V drops down to 15% whereas the pairwise identities across the whole library drops down to 6.52% (**Table 1**). Thus, the library captures the variation around TEV S219V, as well as a considerable evolutionary breadth of the TEV protease family.

To quantify the sequence diversity in the library, we computed the Levenshtein distance between the TEV S219V sequence and each protease sequence in the library (natural homologs and shrunken sequences) (**Fig. 1B**). Within the distribution, there are 3 distinct clusters spanning the approximate ranges from 0-50, 100-150 and 175-225. The 0-50 cluster represents the shrunken TEV sequence design space, with the other two more distant clusters primarily composed of TEV homologs. The presence of distinct clusters rather than a continuous distribution is consistent with underlying clades observed in the phylogenetic tree. Notably, the population of protease homologs with Levenshtein distances less than 100 is very sparse, indicating our homolog library draws from sequences primarily dissimilar from TEV protease. Taken together, these results indicate that the library is highly sequence-diverse.

### Protease Function Persists Across Diverse Genotypes

To examine the relationship between protease function and sequence divergence from TEV S219V, we plotted function measurements against the Levenshtein distance computed for each variant of our dataset (**Fig. 2A**). We defined moderate and high function thresholds in our analysis. The log_10_(intact DHFR) < 3 (moderate function) threshold was selected based on the fitted calibration curve, as values above this point lie outside the curve’s linear regime and thus are close to the assay’s lower-function limit of detection. A more stringent threshold was applied for high function variants (log_10_(intact DHFR) < 2.7), corresponding to comparable activity to TEV S219V. We found that despite high sequence dissimilarity, certain proteases still exhibited high function against the TEV protease substrate. Above the moderate function threshold, sequence alignments revealed a diverse phylogenetic space comprising 66 homologs (**Supplementary File 1**). Since our dataset is compiled from evolutionarily related proteases, we expected functional activity to be preserved in at least some of the TEV homologs under similar selective pressure conditions. However, since substrate specificities diverge across this evolutionary lineage, we would not expect all variants to retain activity on the canonical TEV substrate.^24–26^ For example, TEV and tobacco vein mottling virus proteases share 55% sequence identity, yet catalyze distinct recognition sites with high specificity.^27^ Additional studies examining alternative substrate sequences of homologs throughout the library could elucidate the mechanisms that control recognition and cleavage.

**Figure 2.**
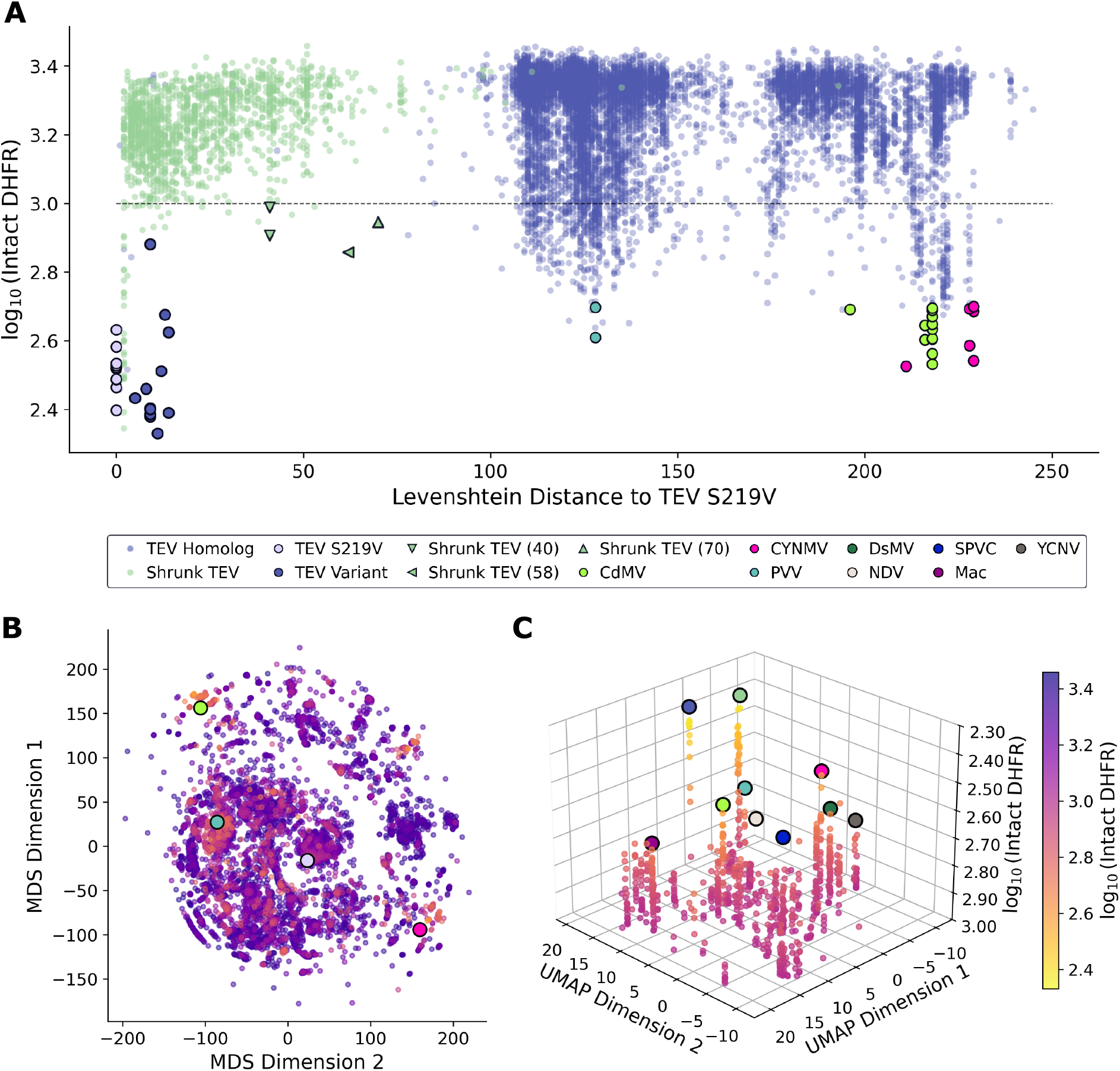
Functional Landscape Across Global and Local Sequence Space **A.** Function measurements plotted against Levenshtein distance to TEV protease. The moderate function threshold is defined here as intact log_10_(intact DHFR) < 3 (black dashed line). Homologs with function comparable to TEV S219V protease (log_10_(intact DHFR) < 2.7) are marked and include cardamom mosaic virus (CdMV), Chinese yam necrotic mosaic virus (CYNMV), and passionfruit Vietnam virus (PVV). Shrunken TEV variants with ≥ 40 deletions are marked by the number of deletions relative to TEV S219V. **B**. 2-dimensional scatter plot of MDS coordinates representing sequence divergence in the global distance landscape. Color indicates protease function, with higher (violet) intact DHFR values representing lower function and lower (yellow) values representing higher function. **C**. 3-dimensional plot showing function plotted against UMAP coordinates representing local distances of similar sequences. The top 10 subgroups ranked by maximum functional variants are shown as large circles and include dasheen mosaic virus (DsMV), sweet potato virus C (SPVC), necrotic degeneration virus (NDV), Macluravirus (Mac), and yam chlorotic necrosis virus (YCNV).

Among the most functional variants in the 175-225 Levenshtein distance range are multiple variants of three proteases from the *Macluravirus* genus: the yam chlorotic necrosis virus, Chinese yam necrotic mosaic virus (CYNMV), and the cardamom mosaic virus (CdMV) proteases. Notably, these proteases differ phylogenically from TEV, which is of the *Potyvirus* genus, yet we observed CYNMV and CdMV function is within the same functional range as TEV protease S219V in this assay.

CdMV and CYNMV variants which exhibited relatively high function against the TEV protease substrate shared just 19% pairwise sequence identity based on BLOSUM62 global alignment. Independent studies have found proteolytic motifs in CdMV and CYNMV that have features similar to those of TEV protease. Jebasingh et. al used TEV protease as a basis to build a CdMV protease structure, and identified a similar catalytic triad (H42, D74, C141) to TEV protease (H46, D81, C151).^28^ Similarly, CYNMV is reported to recognize a similar cleavage site (LQ/M) to TEV protease.^25^

We used AlphaFold 3 to predict the structure of CYNMV, which identified a core structure with high confidence predictions, and a low-confidence N-terminus (**Supplementary Figure 3A**).^29^ Structural alignment using Matchmaker in ChimeraX returned a sequence alignment score of 208.1 and an RMSD = 0.87 *Å* for 93 pruned atom pairs at the core of confidently predicted residues (across all 179 pairs: 7.888) (**Supplementary Fig. 3B**).^30^ This structural alignment suggests that proteolytic function may be robustly preserved through structure despite extreme sequence divergence.

Our dataset includes TEV homologs from databases as well as AI-generated shrunken TEV proteases. In this dataset, we identified TEV protease variants with as many as 14 substitutions with comparable or greater function than TEV S219V. In addition, while most (32 out of 47) shrunken TEV variants with moderate functional activity contained 1 deletion, we identified 4 specific mutants with between 40 and 70 amino acid deletions, retaining activity below our moderate function threshold (represented by green triangles in **Fig. 2A**). Together, these findings illustrate that evolutionarily distant and significantly shrunken sequences can retain proteolytic activity against the TEV substrate.

To explore functional changes in the context of global pairwise distances between sequences, we created a pairwise Levenshtein distance matrix between amino acid sequences and calculated multidimensional scaling (MDS) coordinates using the SMACOF method (**Fig. 2B**). The resulting 2-dimensional plot provides a visualization of sequence divergence across the dataset, where similar sequences are positioned closer together. Overall, we observed high coverage across the MDS landscape, indicating that our library covers a broad range of sequence diversity. Functional variants are dispersed throughout the embedding, indicating function persists across diverse genotypes. For example, notable homologs with strong function, such as CYNMV, CdMV, and PVV occupy spatially distinct regions. This builds on the observation in **Fig. 2A** that sequences dissimilar to TEV S219V can remain functional. These functional variants are not only highly divergent from TEV S219V but also from one another, with a pairwise identity of 46.67% between CdMV and CYNMV. The appearance of functional variants through the global MDS space suggests sequence-flexibility for preserving function.

Since MDS prioritizes global distances between clusters of sequences, we wanted to investigate local sequence landscapes to identify closely related variants. We computed uniform manifold approximation and projection (UMAP) embeddings, which preserve local, neighborhood structure, and visualized functional variants in 3-dimensions (**Fig. 2C**). We applied a function threshold (<3) to highlight closely related variants with at least moderate function, and identified clusters within the UMAP space using DBSCAN.^31^ We ranked the clusters according to local function minima, and highlighted the top 10 of these clusters to identify groups of high-functioning homologs within the dataset that share similar sequences. When grouped by local distances, similar sequences display a broad range of function, indicating sensitivity to mutation that can be explored in future assays.

## Conclusion

We characterized a library of 11,722 TEV protease homologs and AI-generated variants to explore a highly diverse sequence-function landscape. This library has a minimum sequence identity of 6.52% and a mean of 41.79%. We measured the function of all variants using our GROQ-seq assay, which links split-DHFR substrate cleavage to cellular fitness. Despite sequence divergence, we identified several evolutionary distinct protease homologs with comparable function to TEV protease throughout the global distance landscape. In one example, alignment of the predicted CYNMV structure with TEV protease revealed similar structural motifs despite 19% pairwise sequence identity. When grouped by local distances, similar sequences display a broad range of function. Remarkably, we also identified TEV protease variants with as many as 14 substitutions with comparable function to TEV S219V. Additionally, we observed shrunken TEV proteases with as many as 70 deletions that still cleave the TEV protease substrate. These heavily-mutated TEV protease variants within our dataset demonstrate that the TEV protease sequence can be significantly varied yet still preserve proteolytic activity. These findings demonstrate that future BMS studies can apply the GROQ-seq assay to explore additional functions with sequence-diverse libraries.

Our results show that despite substantial phylogenetic divergence across genera, many naturally occurring proteases retain the ability to recognize and cleave the same sequence targeted by TEV protease. Future work could investigate the features of the recognition site and catalytic residues to improve sequence-structure–function predictive models. These proteases can also be tested against varied substrate sequences to further elucidate the interplay between protease structure and substrate specificity.

In summary, we employed BMS via GROQ-seq to produce a large dataset of AI-ready biological data, quantifying protease function. This dataset contributes to the need for large, sequence-diverse training data and may help reduce bias in machine learning models.

## Supplementary Information

**Supplementary Figure 1.**
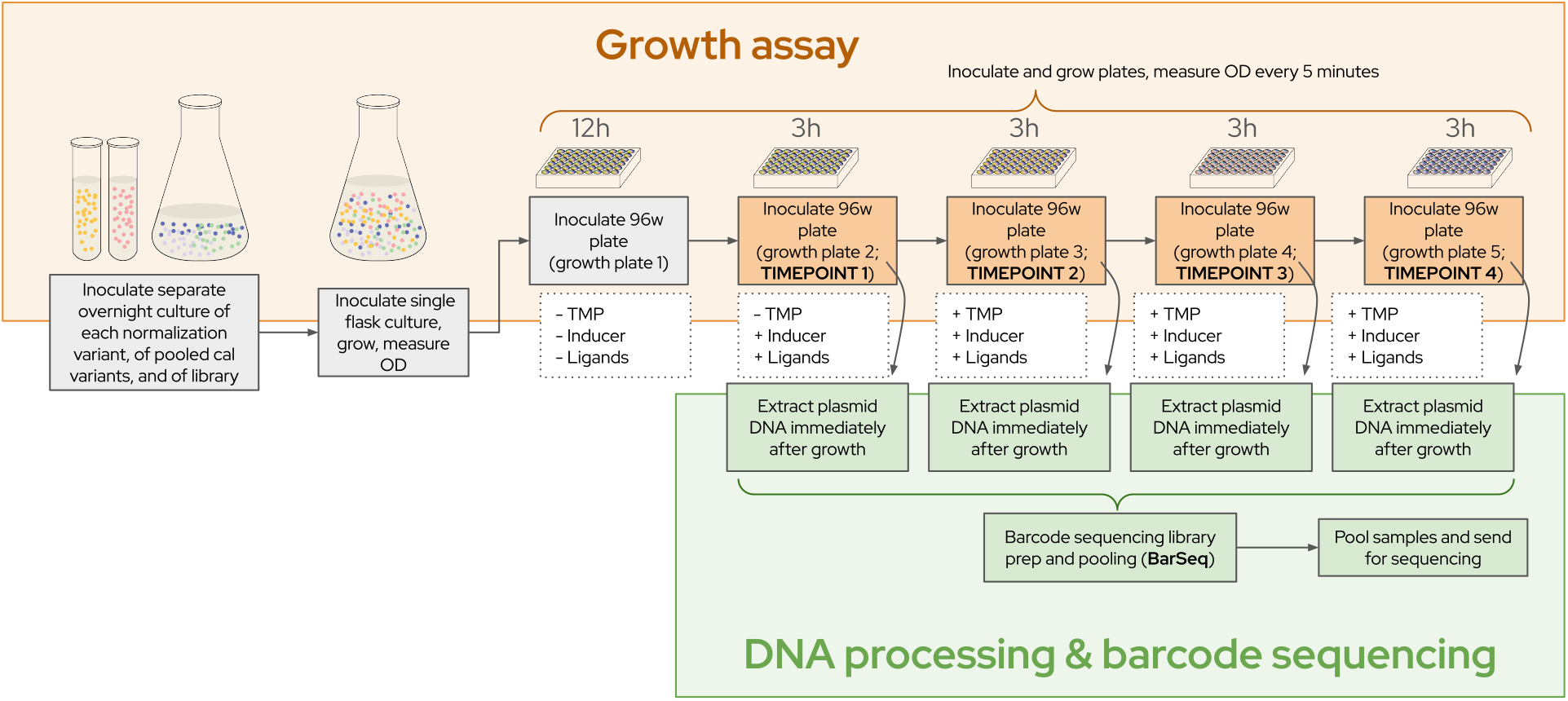
Schematic representation of the GROQ-seq workflow

**Supplementary Figure 2.**
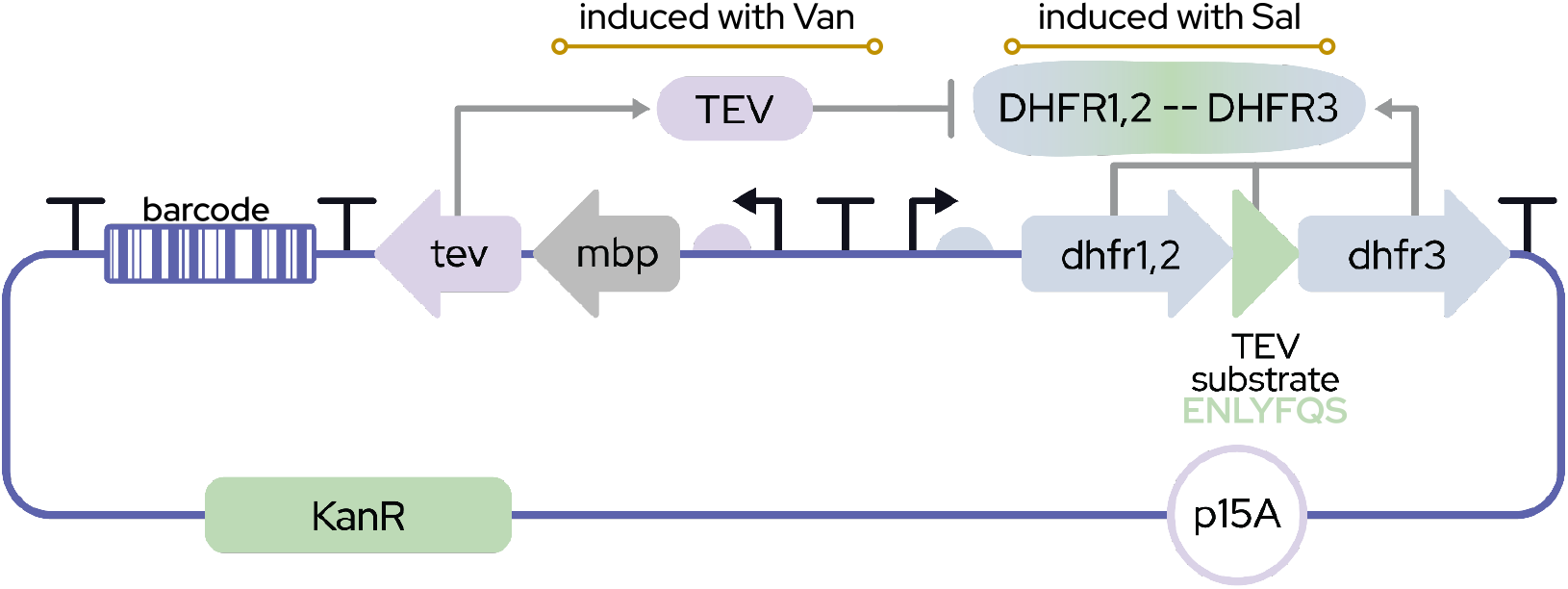
Schematic representation of the genetic circuit used in this system

**Supplementary Figure 3.**
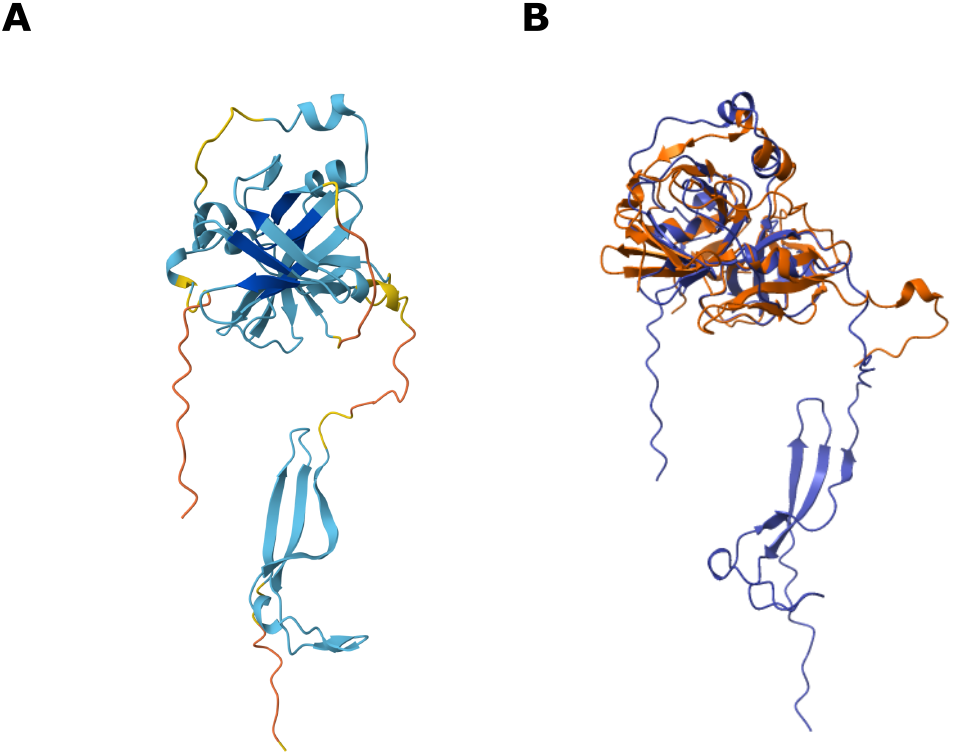
CYNMV predicted structure comparison to TEV protease structure **A**. AlphaFold 3 structural predictions of Chinese yam necrotic mosaic virus (CYNMV) colored by computed pIDDT. Dark blue: pIDDT >90. Light blue: 90 > pIDDT > 70. Yellow: 70 > pIDDT > 50. Orange: pIDDT < 50. **B**. Structural alignment of CYNMV predicted structure (purple) and TEV Protease (1LVM) (orange).

**Supplementary Table 1.**
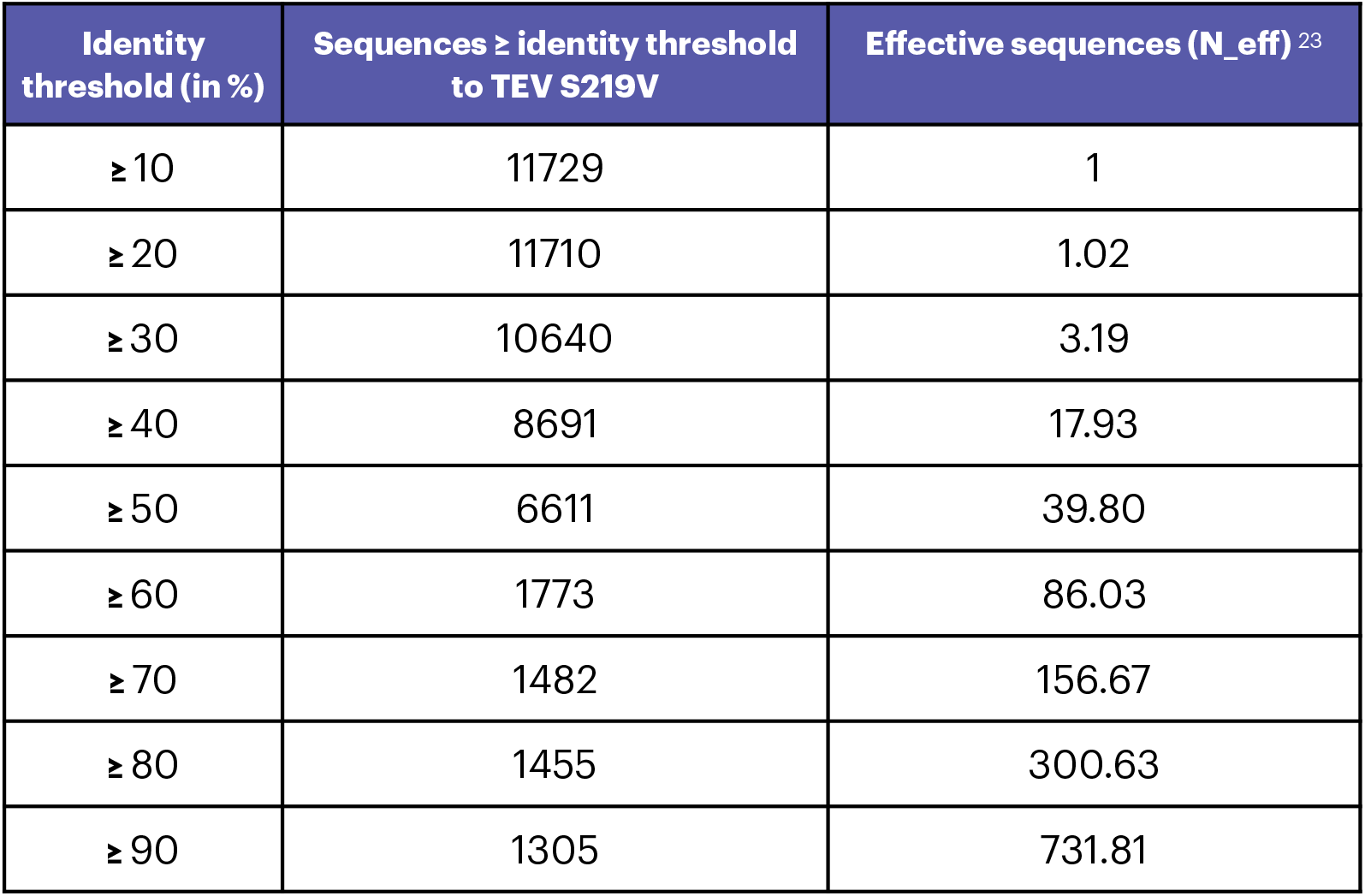
Library composition across different pairwise sequence identity thresholds. These values were computed from the MAFFT multiple sequence alignment restricted to 233 match columns with <50% gaps across all sequences)

## Funder Information Declared

The Align Foundation is supported by Griffin Catalyst and Schmidt Sciences.

## Disclaimer

Certain commercial equipment, instruments, or materials are identified in this paper in order to specify the experimental procedure adequately. Such identification is not intended to imply recommendation or endorsement by the National Institute of Standards and Technology, nor is it intended to imply that the materials or equipment identified are necessarily the best available for the purpose.

